# HLH-30 dependent rewiring of metabolism during starvation in *C. elegans*

**DOI:** 10.1101/2020.06.26.170555

**Authors:** Kathrine B. Dall, Jesper F. Havelund, Eva Bang Harvald, Michael Witting, Nils J. Færgeman

**Author notes:** To whom correspondence should be addressed: Nils J. Færgeman.

## Abstract

One of the most fundamental challenges for all living organisms is to sense and respond to alternating nutritional conditions in order to adapt their metabolism and physiology to promote survival and achieve balanced growth. Here, we applied metabolomics and lipidomics to examine temporal regulation of metabolism during starvation in wildtype *Caenorhabditis elegans* and in animals lacking the transcription factor HLH-30. Our findings show for the first time that starvation alters the abundance of hundreds of metabolites and lipid species in a temporal- and HLH-30-dependent manner. We demonstrate that premature death of *hlh-30* animals under starvation can be prevented by supplementation of exogenous fatty acids, and that HLH-30 is required for complete oxidation of long-chain fatty acids. We further show that RNAi-mediated knockdown of the gene encoding carnitine palmitoyl transferase I (*cpt-1*) only impairs survival of wildtype animals and not of *hlh-30* animals. Strikingly, we also find that compromised generation of peroxisomes by *prx-5* knockdown renders *hlh-30* animals hypersensitive to starvation, which cannot be rescued by supplementation of exogenous fatty acids. Collectively, our observations show that mitochondrial functions are compromised in *hlh-30* animals and that *hlh-30* animals rewire their metabolism to largely depend on functional peroxisomes during starvation, underlining the importance of metabolic plasticity to maintain survival.

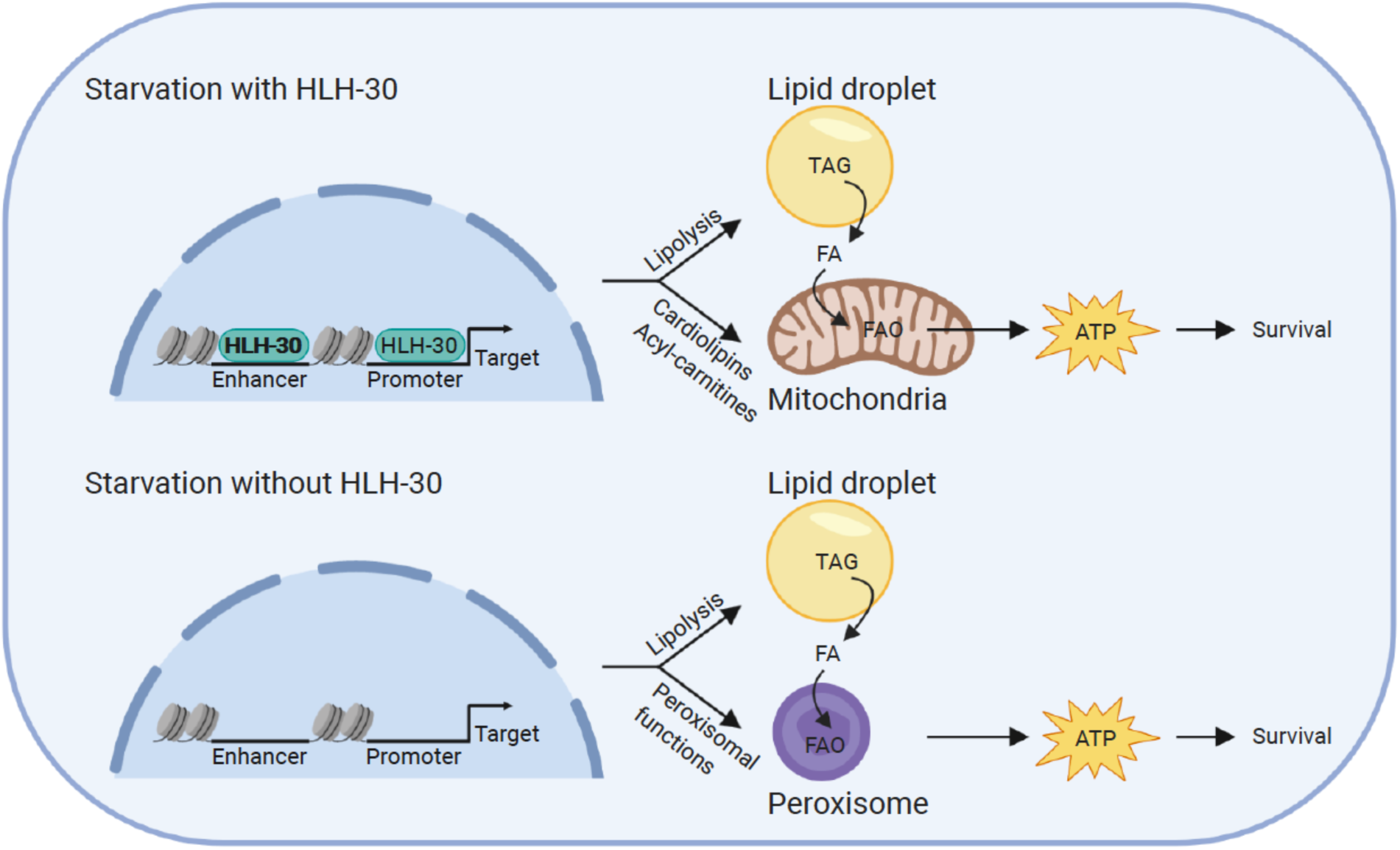

## 1 INTRODUCTION

The ability to regulate metabolism in response to changes in nutrient availability is an evolutionarily conserved mechanism ranging from bacteria to humans. Regulating metabolism by coordinating anabolic and catabolic pathways to sustain metabolic homeostasis ensures prolonged survival during periods of nutrient scarcity. However, rewiring of energy metabolism is a complex and dynamic process encompassing many transcriptional and post-transcriptional regulators. One of the fundamental mechanisms promoting survival in response to starvation is the use of energy stores e.g. via breakdown of lipids. Fatty acids are mobilized from lipid droplets in adipocytes for mitochondrial β-oxidation through either lipolysis or lipophagy, pathways that have both been reported to be crucial for surviving starvation (Martin & Parton, 2006). A major regulator of lipid metabolism during starvation is the conserved basic helix-loop-helix transcription factor HLH-30 in *Caenorhabditis elegans* (*C. elegans*), an ortholog of the mammalian transcription factor EB (TFEB) (Lapierre et al., 2013). HLH-30/TFEB regulates the expression of genes belonging to the Coordinated Lysosomal Expression And Regulation (CLEAR) network, which are involved in autophagosome formation, lysosomal biogenesis, lipase function, and fatty acid degradation (Martina et al., 2014; Palmieri et al., 2011; Settembre et al., 2013). HLH-30/TFEB is also a transcription factor mediating resistance to several stressors besides starvation, including oxidative stress, heat stress, and host defense against pathogen infection (Lin et al., 2018; Visvikis et al., 2014). Additionally, removal of HLH-30/TFEB impairs the longevity of several long-lived *C. elegans* mutants (Lapierre et al., 2013) and the *hlh-30* mutant itself dies prematurely during starvation (Harvald et al., 2017; O’Rourke & Ruvkun, 2013). In the present study, we have successfully applied a combinatorial metabolomics and lipidomics approach to examine temporal regulation of metabolism during starvation and how HLH-30 regulates metabolism during starvation in *C. elegans*. Specifically, we find that starvation induces significant and specific changes in the metabolome and in the lipidome of *C. elegans*. In particular, we find that starvation induces both long-chain acyl-carnitine and cardiolipin levels in wildtype animals, in accordance with enhanced mitochondrial metabolism. Accordingly, we find that starvation induces oxidation of oleic acid. Markedly, induction of cardiolipin and acyl-carnitine levels as well as oxidation of oleic acid upon starvation are completely absent in *hlh-30* animals, arguing that HLH-30 is required for induction of mitochondrial β-oxidation during starvation. Interestingly, we find that impaired generation of peroxisomes induces premature death of *hlh-30* animals upon starvation, which cannot be rescued by supplementation of exogenous fatty acids. Collectively, we show for the first time, that functional loss of HLH-30 renders *C. elegans* highly dependent on peroxisomal degradation of fatty acids to survive starvation. Our observations substantiate the importance of metabolic plasticity in order to survive periods of nutrient scarcity.

## 2 RESULTS

### Starvation induced metabolic and lipidomic re-arrangement enhances mitochondrial function in C. elegans

To identify the metabolic response to starvation we analyzed the temporal response to starvation in *C. elegans* across a 16 hours starvation time course at the mid-L4 stage by metabolomics and lipidomics (Figure 1). As previously (Harvald et al., 2017), we analyzed multiple time points within the first 6 hours to interrogate the early starvation responses with high resolution. We harvested animals in biological triplicate at each of the 7 time points for extraction of metabolites and lipids, respectively, and for subsequent analyses by MS-based metabolomics and lipidomics (Figure 1). To optimize lipid extraction from *C. elegans*, we applied different commonly used lipid extraction methods and performed lipid profiling in positive ionization mode. By using the BUME extraction (Lofgren et al., 2012) we detected 4.963 different molecular features in the apolar phase, while we detected 4819, 4249, 5172, 5082 molecular features when using Bligh and Dyer (Bligh & Dyer, 1959), Folch (Folch, Lees, & Sloane Stanley, 1957), MMC (Pellegrino, Di Veroli, Valeri, Goracci, & Cruciani, 2014), and MTBE (Matyash, Liebisch, Kurzchalia, Shevchenko, & Schwudke, 2008) extraction methods, respectively (Figure S1). 4143 features were commonly detected in all tested extraction methods (Figure S1). Although, the MTBE extraction yielded the most features, we also found that this method showed the highest extraction variability. In contrast, the Folch extraction showed the lowest variability, but also the lowest number of features (Figure S1). Based on the overall performance, the BUME extraction method not only showed a high number of features with a low variability, but also provided a polar phase for LC-MS metabolomic analyses of polar metabolites. Furthermore, it showed better recovery of polar lipids such as lysolipids compared to the other methods. Applying the BUME extraction and lipid profiling to our samples, 4063 lipid features in positive and 2258 in negative ionization mode remained after normalization and filtering (detected in all QCs and RSD < 30%). Out of these, 2068 were putatively annotated on the MS^1^ level and 427 on the MS^2^ level in positive ionization mode, 955 and 118 in negative mode, respectively (Table S1). We generated volcano plots to visualize to which extend starvation rewires the *C. elegans* metabolome across a 16 hours starvation time course (Figure 2a and Figure S2). This showed that short-term starvation (1 and 2 hours) only has subtle effects on the metabolome in wildtype animals, while starvation for 3 hours profoundly changes the metabolome (61 significantly altered metabolites in total in negative and positive modes) (Table S1). Further, we found that the abundance of 289, 271 and 501 molecular features was significantly altered after 4, 6, and 16 hours of starvation, respectively (Figure 2a, Figure S2 and Table S1). During starvation, lipolysis of triacylglycerols plays an important role as an energy source in both mammals and in *C. elegans* (Buis et al., 2019; Lee et al., 2014; Martin & Parton, 2006; Murphy et al., 2019; Zaarur et al., 2019). Accordingly, the fatty acids released by lipolysis are subsequently activated to CoA-esters and transported into the mitochondria by the carnitine-shuttle system for degradation by β-oxidation. Consistently, we observed that the level of long-chain acyl-carnitines massively increased already after one hour and remained elevated after 16 hours of starvation in wildtype animals, while levels of short-chain acyl-carnitines largely remained unchanged (Figure 2a and Figure 3a).

**Figure 1.**
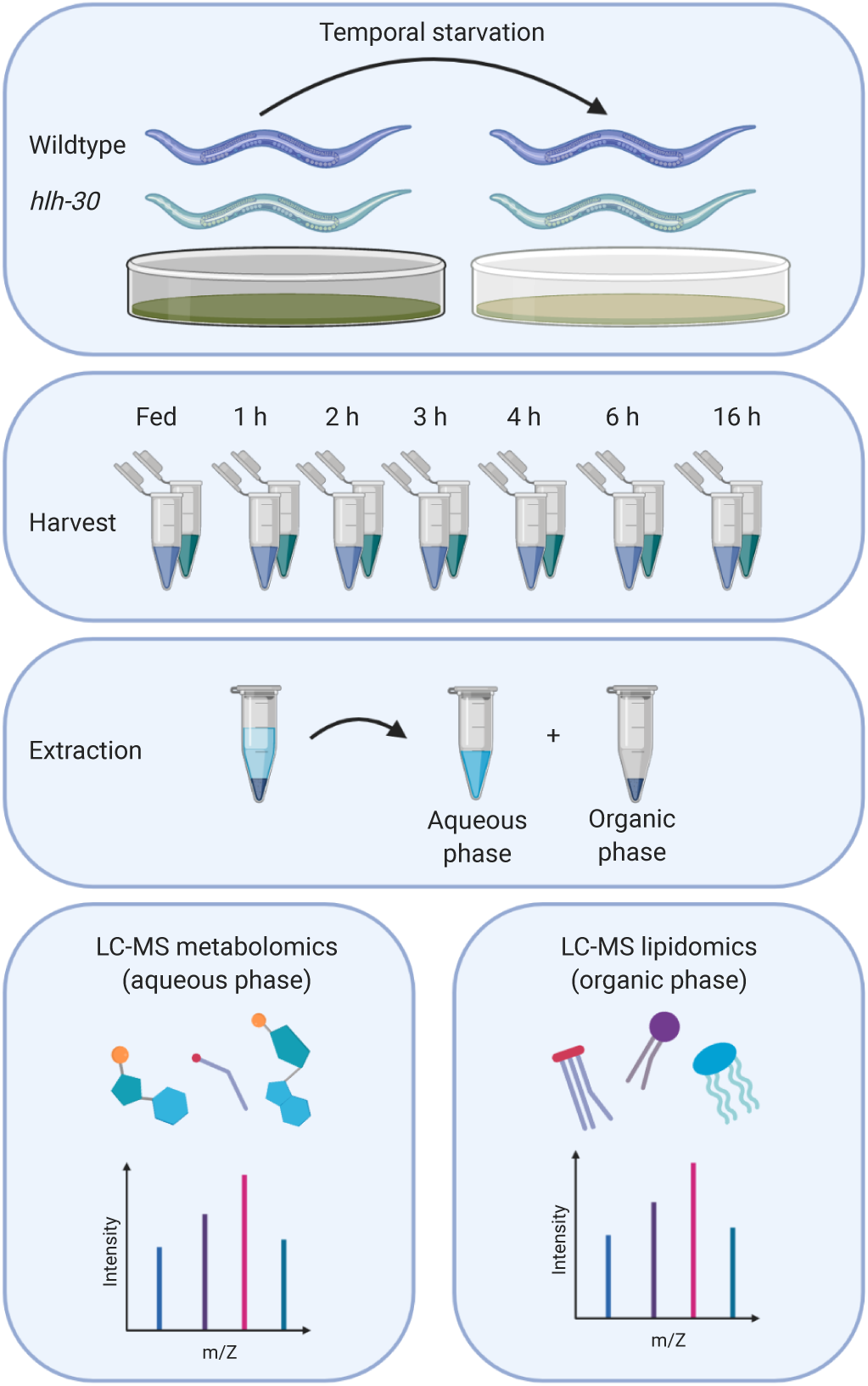
Experimental workflow for combined metabolomics and lipidomics of the starvation response in *C. elegans*. Wildtype *C. elegans* and the *hlh-30* mutant were included in the study. Starvation was induced by transferring animals to plates containing no bacteria. Worms were starved for 1, 2, 3, 4, 6 and 16 hours of starvation, prior to harvesting and extraction of metabolites and lipids. The upper aqueous phase was analyzed using LC-MS based metabolomics and the lower organic phase was analyzed using LC-MS based lipidomics.

**Figure 2.**
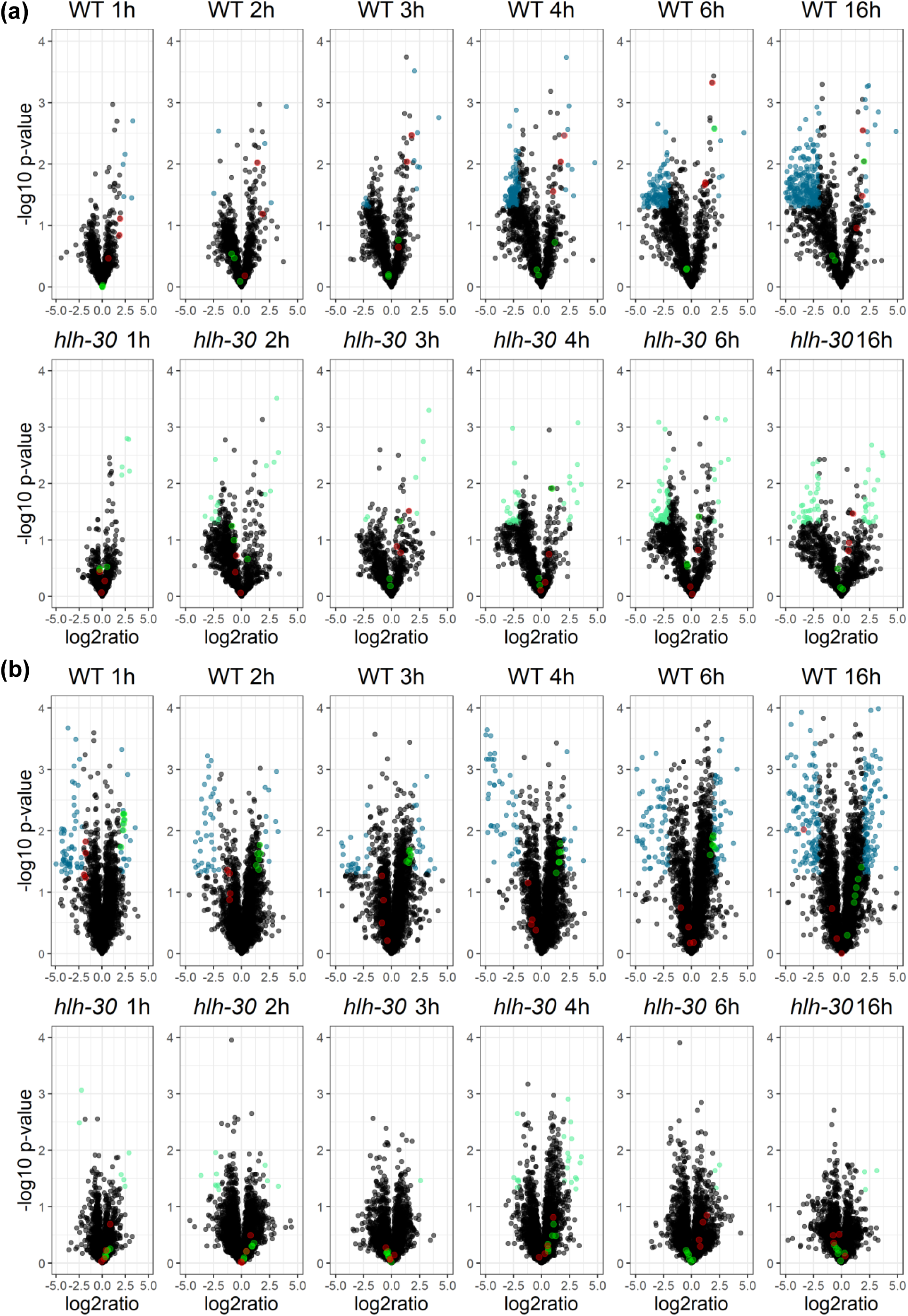
Metabolic and lipidomic changes induced by temporal starvation. (a) Volcano plot displaying changes in the metabolome in response to starvation for the wildtype and *hlh-30* mutant at each timepoint. Significant up- or down-regulated metabolites in wildtype animals are shown in blue, and in light green for the *hlh-30* animals. Regulation of long-chain acyl-carnitines are shown in red and short-chain acyl-carnitines are shown in green for both wildtype and the *hlh-30* mutant. Only metabolites detected in the positive mode are shown. (b) Volcano plot displaying changes in the lipidome in response to starvation for the wildtype and *hlh-30* animals at each timepoint. Significant up- or down-regulated lipids in the wildtype are shown in blue, and in light green for *hlh-30* animals. Regulation of cardiolipins are shown in green and N-acylethanolamines are shown in red for both wildtype and *hlh-30* animals. Only lipid species detected in the positive mode are shown. Metabolites and lipids with a *p*-value < 0.05 and a fold-change of > 2 or < 0.5 were considered to be significantly changed.

**Figure 3.**
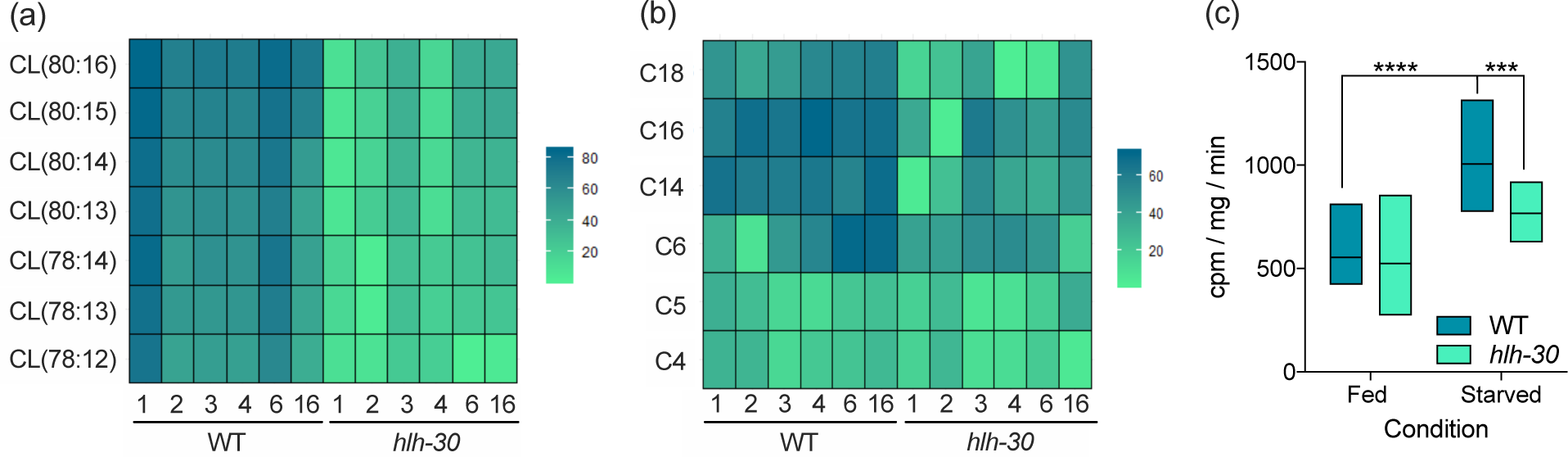
Acyl-carnitine and cardiolipin regulation is disrupted in the *hlh-30* mutant. (a) Heatmap illustrating the log_2_ fold-change of specific long-chain acyl-carnitines at each starvation timepoint compared to fed condition for the wildtype and *hlh-30* mutant. (b) Heatmap illustrating the log_2_ fold-change of specific cardiolipins at each starvation timepoint compared to fed condition for the wildtype and *hlh-30* mutant. Blue: upregulation, green: downregulation. (c) Oxidation of ^3^H-labeled oleic acid is shown for wildtype and *hlh-30* animals in fed state and after 6 hours of starvation ****: *p*-value < 0.0001, ***: *p*-value < 0.0009.

By similar means, we visualized how the *C. elegans* lipidome alters across the starvation time course (Figure 2b and Figure S3). The volcano plots clearly show that starvation also rewires the lipidome already after 1 hour, as 79 and 99 lipid species were significantly up- or down regulated in positive and negative mode, respectively (Figure 2b, Figure S3 and Table S1). Consistent with previous observations (Lucanic et al., 2011), we also found that the abundance of lipids like N-acylethanolamines (NAEs) decreased in wildtype animals across the starvation time course (Figure 2b). Changed lipids were grouped according to their profile across time. Based on this, lipids could be separated into species changing early within the first time points and later responders, e.g. triacylglycerols show changes at 16 hours of starvation. Moreover, among the upregulated lipid species we identified seven lipid species in wildtype animals that significantly increased across all time points expect after 16 hours of starvation. Based on their elution profile and MS-fragmentation pattern, we identified these lipid species to be cardiolipins (Figure S4). Cardiolipins are major phospholipids almost exclusively located in the inner mitochondrial membrane and required for mitochondrial morphology, mitochondrial membrane dynamics and energy production not only in *C. elegans* but also in other eukaryotes (Sakamoto et al., 2012; Sustarsic et al., 2018). Collectively, these results show that starvation induces major re-arrangements of both the metabolome and lipidome, and that mitochondrial functions are enhanced by starvation in wildtype *C. elegans*.

### Rewiring of lipid metabolism and induction of β-oxidation upon starvation depend on HLH-30 in C. elegans

Since the transcription factor HLH-30 and its mammalian ortholog TFEB previously have been shown to serve crucial functions during starvation, dietary restriction, and autophagy in both *C. elegans* and in mammals (Harvald et al., 2017; Lapierre et al., 2013; Murphy et al., 2019; O’Rourke & Ruvkun, 2013; Roczniak-Ferguson et al., 2012; Settembre et al., 2013) this prompted us to examine how functional loss of HLH-30 modulates the metabolome as well as the lipidome in response to starvation in *C. elegans*. In contrast to wildtype animals, we only found a limited number of metabolites and lipid species that changed significantly in response to starvation in *hlh-30* animals (Figure 2b and Table S1). Markedly, we found that the level of long-chain acyl-carnitines in *hlh-30* animals remained largely unchanged in response to starvation compared to wildtype animals (Figure 2a and Figure 3a), arguing that fatty acid import into mitochondria is compromised. In keeping with this notion, we also found that cardiolipin levels in *hlh-30* animals largely remained unchanged in response to starvation (Figure 2b and Figure 3b). We therefore speculate that HLH-30/TFEB is required for biogenesis or for the maintenance of functional mitochondria. Thus, to corroborate these observations, we assessed how functional loss of HLH-30 affected fatty acid oxidation during starvation in *C. elegans*, by examining complete oxidation of ^3^H-labeled oleic acid. During fed conditions, oxidation of oleic acid is similar in *hlh-30* animals and in wildtype animals. However, consistent with previous observations, oxidation of oleic acid increased significantly after six hours of starvation in wildtype animals, while it remained unchanged in *hlh-30* animals (Figure 3c). This observation substantiates that HLH-30 is required for induction of fatty acid oxidation during starvation and hence for metabolic adaptation during starvation.

### Fatty acid supplementation rescues premature death during starvation in the hlh-30 mutant

TFEB, the mammalian ortholog of HLH-30, has recently been found to be required for mitochondrial biogenesis, morphology, and functions in skeletal muscle in mice (Mansueto et al., 2017). Although loss of HLH-30 functions does not affect mitochondria morphology in *C. elegans* (Murphy et al., 2019), the present observations show that HLH-30 is required to support fundamental mitochondrial functions in *C. elegans* during limited nutritional conditions. Compared to wildtype animals, *hlh-30* animals die prematurely during starvation (Harvald et al., 2017). Since mobilization of fatty acids from intestinal lipid stores is required for *C. elegans* to withstand long-term starvation (Buis et al., 2019), we therefore hypothesized that exogenous supplementation of medium-chain fatty acids, which cross the mitochondrial membranes independent of the carnitine-shuttle system, would rescue the premature death of *hlh-30* animals. We therefore examined survival under starvation conditions by transferring animals to empty plates supplemented with either a medium-chain (lauric acid, C_12:0_) or a long-chain fatty acid (palmitic acid, C_16:0_). As previously, we found that *hlh-30* animals die prematurely during starvation when compared to wildtype animals. However, when supplemented with lauric acid both wildtype and *hlh-30* animals survived significantly longer during starvation when compared to un-supplemented animals (Figure 4 and Table S2). In fact, *hlh-30* animals were completely rescued to wildtype levels by lauric acid. Supplementation with palmitic acid also extended the survival of wildtype animals and surprisingly also of *hlh-30* animals, however not to the same extend as lauric acid (Figure 4 and Table S2).

**Figure 4.**
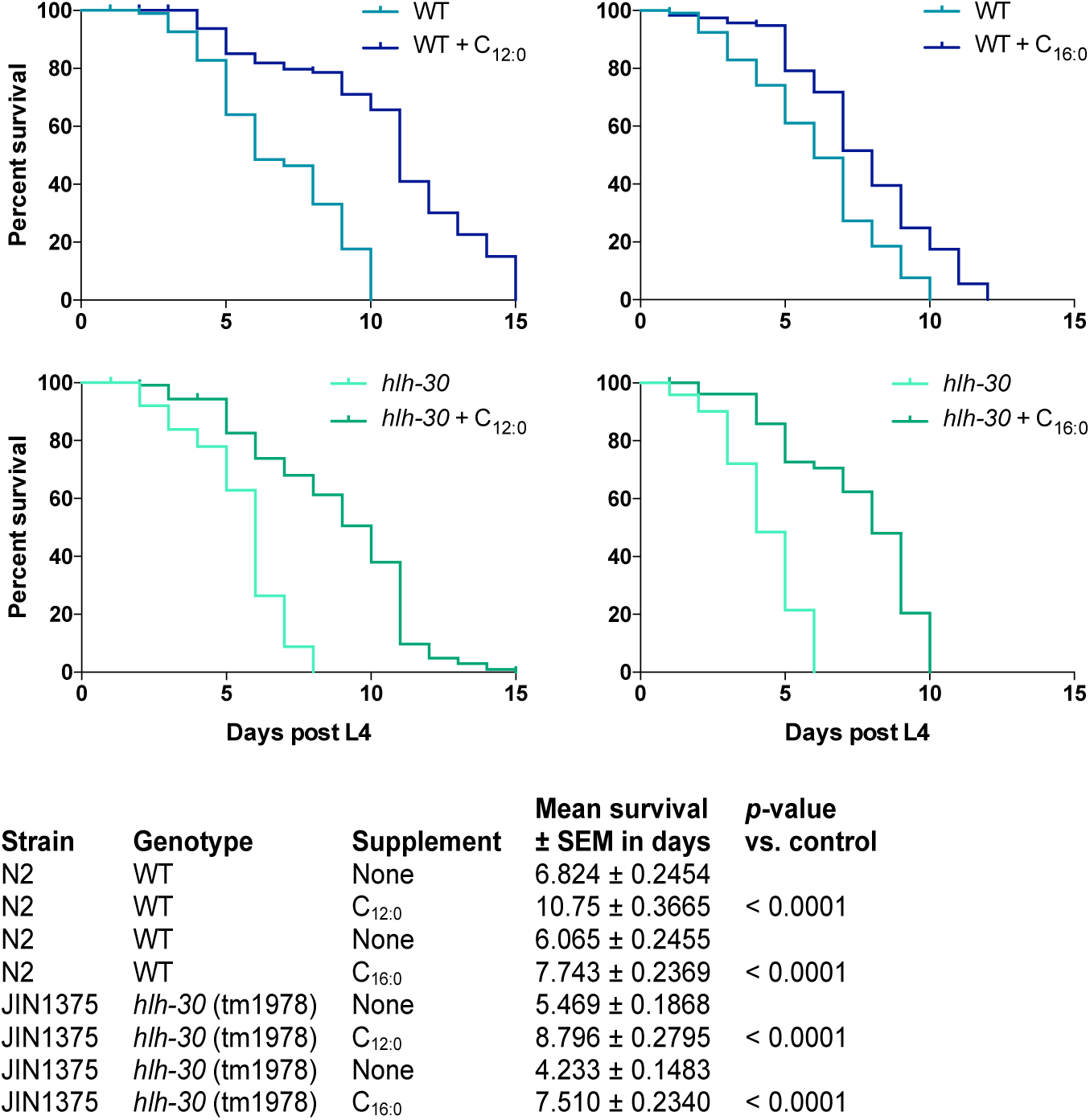
Exogenous fatty acid supplementation rescues premature death of the *hlh-30* mutant during starvation. Survival-span showing the effect of fatty acid supplementation on both wildtype (blue) and *hlh-30* mutant (green) survival in response to starvation. Starvation was induced at start L4 stage by transferring worms to empty plates. Both strains were supplemented with either vehicle, a medium-chain fatty acid, lauric acid (C_12:0_) or a long-chain fatty acid, palmitic acid (C_16:0_) throughout the experiment. Survival was monitored every day. Survival analysis was carried out using the Kaplan-Meier estimator and *p*-value was calculated using log-rank test in GraphPad Prism 6.

### Disruption of the carnitine-shuttle system impairs starvation survival of wildtype animals

Our findings support the notion that impaired β-oxidation, caused by e.g. diminished mitochondrial import of long-chain fatty acids, may be the underlying reason for the inability of the *hlh-30* mutant to survive during starvation. By RNA-sequencing Harvald et al. recently profiled the genome-wide response to starvation (Harvald et al., 2017), and found that expression of genes encoding lipases needed for conventional lipolysis of triacylglycerols in lipid droplets (*atgl-1*) or via lysosomal breakdown (*lipl-2 to lipl-4*) increased in wildtype animals upon starvation but remained constant or decreased in *hlh-30* animals during starvation (Figure S5). Similarly, the expression of genes encoding enzymes required for activation of fatty acids (*acs-2*) and for active transport of fatty acids into the mitochondria (*cpt-1*) is diminished in the mutant upon starvation (Figure S5). We therefore speculated that downregulation of *cpt-1* expression would impair the ability of wildtype animals to survive under starvation conditions. Intriguingly, we found that RNAi-mediated knockdown of *cpt-1* significantly impairs survival of wildtype animals under starving conditions compared to the control animals, while *cpt-1* knockdown had no effect on survival of *hlh-30* animals (Figure 5 and Table S3). Markedly, lauric acid supplementation fully rescued the effects of *cpt-1* knockdown in wildtype animals and extended survival of *hlh-30* animals to wildtype levels independent of *cpt-1* knockdown. Consistent with the notion that mitochondrial import of long-chain fatty acids depends on a functional carnitine-shuttle system, palmitic acid (C_16:0_) supplementation did not fully rescue survival of wildtype control animals. Interestingly, palmitic acid supplementation rescued survival of *hlh-30* animals under starvation conditions independent of *cpt-1* knockdown, indicating that fatty acids can support survival during starvation by being channeled to other energy-producing pathways than mitochondrial β-oxidation.

**Figure 5.**
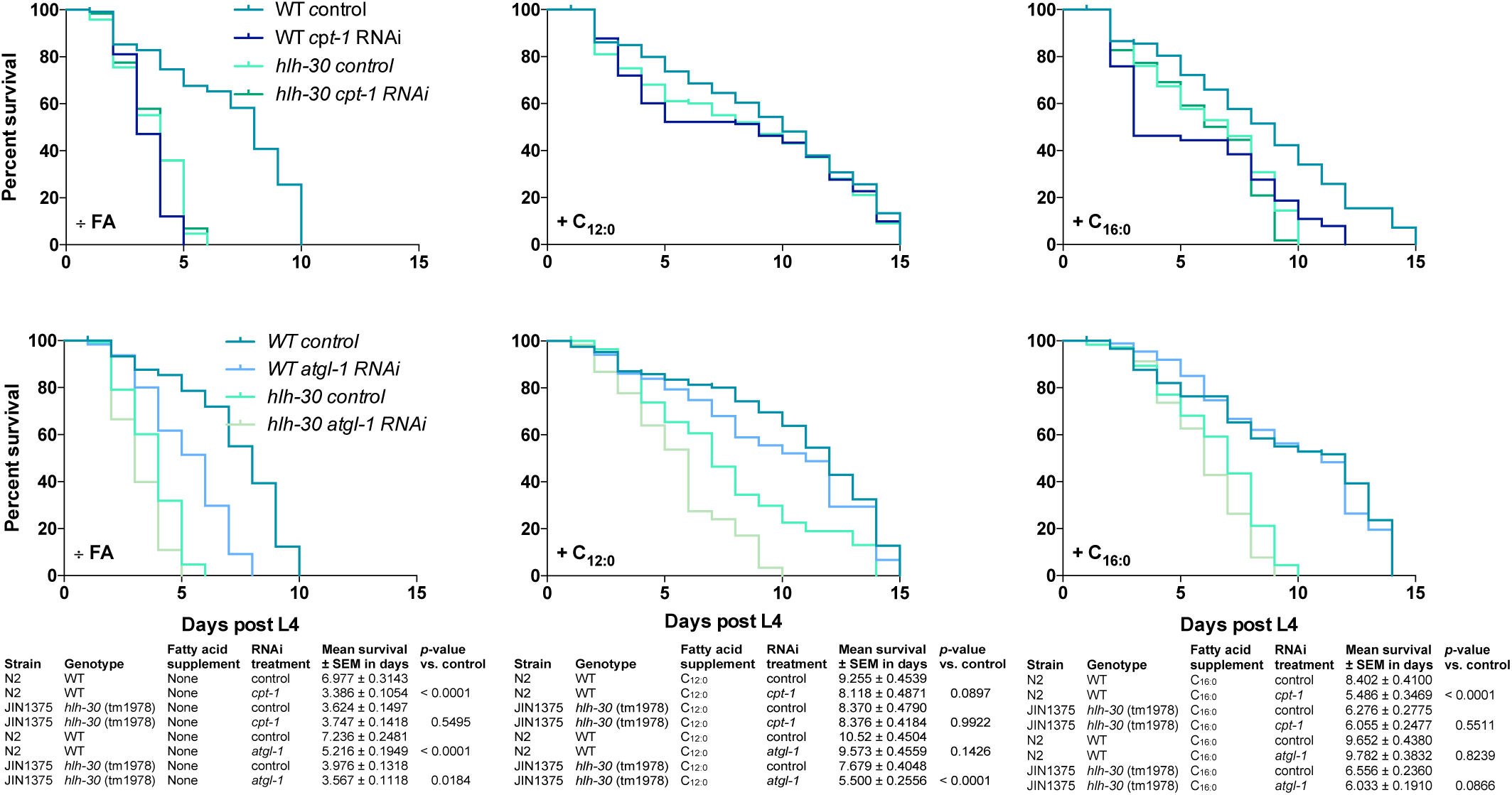
Survival of wildtype animals is impaired by disruption of the carnitine-shuttle. (a) Survival-span showing the effect of RNAi mediated knockdown of *cpt-1* on the survival of wildtype and *hlh-30* to mutant during starvation. Starvation was induced at start L4 stage by transferring worms to empty plates. Both strains were supplemented with either a medium-chain fatty acid, lauric acid (C_12:0_) or a long-chain fatty acid, palmitic acid (C_16:0_) throughout the experiment. Survival was monitored every day. (b) Survival-span showing the effect of RNAi mediated knockdown of *atgl-1* on the survival of wildtype and *hlh-30* to mutant during starvation. Starvation was induced at start L4 stage by transferring worms to empty plates. Both strains were supplemented with either a medium-chain fatty acid, lauric acid (C_12:0_) or a long-chain fatty acid, palmitic acid (C_16:0_) throughout the experiment. Survival was monitored every day. Survival analysis was carried out using the Kaplan-Meier estimator and *p*-value was calculated using log-rank test in GraphPad Prism 6.

ATGL-1 mediates lipolysis of intestinal lipid stores during fasting in *C. elegans* (Lee et al., 2014). We therefore assessed whether the mobilization of stored lipids would affect survival during starvation. Expectedly, knockdown of *atgl-1* in wildtype animals significantly shortened survival compared to its control during starvation but had only minor effects on the survival-span of *hlh-30* animals (Figure 5b). Supplementation with lauric and palmitic acid both extended survival-span of wildtype and *hlh-30* animals, yet only lauric acid extended survival-span to wildtype levels. All together, we interpret these observations that mobilization and mitochondrial import of fatty acids from lipid stores are crucial for surviving during starvation, however only the latter is dependent on HLH-30 in *C. elegans*.

### Survival of the hlh-30 mutant during starvation is dependent on peroxisomal β-oxidation

Since supplementation of palmitic acid also improved survival of *hlh-30* animals during starvation made us speculate whether *hlh-30* animals compensate by using alternative metabolic pathways to generate sufficient energy to survive starvation. Besides mitochondria, peroxisomes are also capable of degrading fatty acids. Like mitochondrial β-oxidation, peroxisomal β-oxidation catalyzes chain shortening of acyl-CoAs by four enzymatic steps yielding acetyl-CoA. Despite that peroxisomal β-oxidation in *C. elegans* is mostly known for oxidizing very long-chain fatty acids (VLCFAs) and for the synthesis of ascarosides (Artyukhin et al., 2018), long- and medium chain saturated and unsaturated fatty acids can also serve as substrates for peroxisomal β-oxidation (Poirier, Antonenkov, Glumoff, & Hiltunen, 2006). The first step is catalyzed by the enzyme acyl-CoA oxidase (ACOX), considered to be the main regulator of the flux through the pathway. Interestingly, expression of *acox* genes in *hlh-30* animals is upregulated compared to wildtype animals and sustain upregulated through starvation (Harvald et al., 2017) (Figure S5). Therefore, since mitochondrial functions are compromised in *hlh-30* animals, we speculated that they rewire their metabolism towards peroxisomes in order to survive starvation. Thus, by RNAi we knocked down *prx-5* that encodes a peroxisomal assembly factor required for biogenesis of functional peroxisomes (Weir et al., 2017). Intriguingly, upon knockdown of *prx-5*, survival of *hlh-30* animals during starvation was dramatically decreased when compared to its control, while *prx-5* knockdown in wildtype animals did not affect survival during starvation (Figure 6 and Table S3). Moreover, neither lauric acid nor palmitic acid could rescue the effect of *prx-5* knockdown, collectively implying that *hlh-30* animals switch their metabolic program towards peroxisomes.

**Figure 6.**
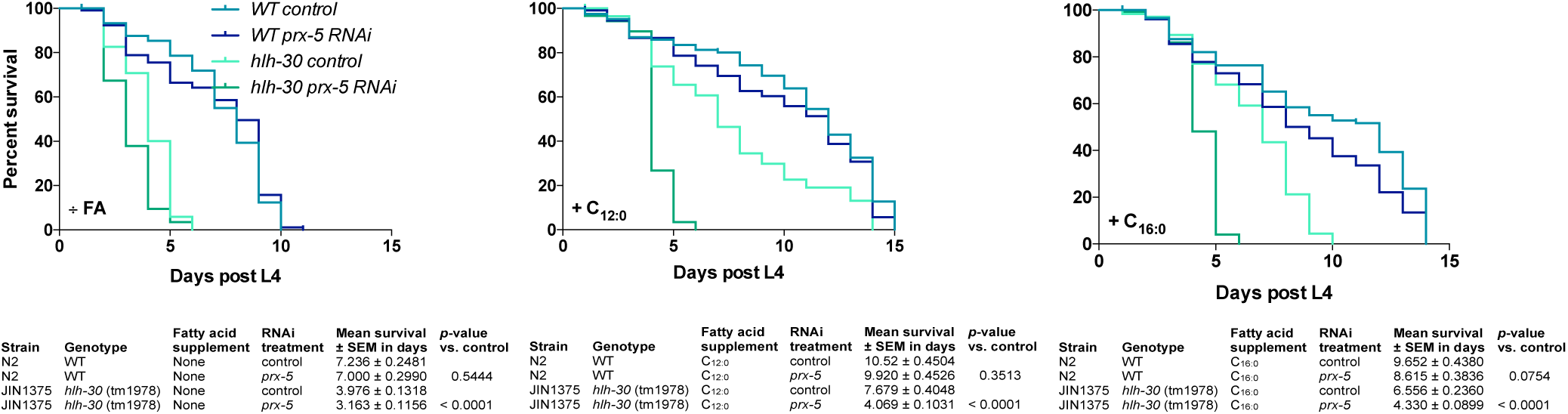
The ability of the *hlh-30* mutant to survive starvation is dependent on peroxisomal β-oxidation. Survival-span showing the effect of RNAi mediated knockdown of *prx-5* on the survival of wildtype and *hlh-30* to mutant during starvation. Starvation was induced at start L4 stage by transferring worms to empty plates. Both strains were supplemented with either a medium-chain fatty acid, lauric acid (C_12:0_) or a long-chain fatty acid, palmitic acid (C_16:0_) throughout the experiment. Survival was monitored every day. Survival analysis was carried out using the Kaplan-Meier estimator and *p*-value was calculated using log-rank test in GraphPad Prism 6.

In conclusion, by using a systems-wide analyses we have demonstrated how loss of the transcription factor HLH-30 globally rewires metabolism in the nematode *C. elegans* during starvation. We found that animals lacking the transcription factor HLH-30 fails to upregulate cardiolipin synthesis and β-oxidation during starvation and compensate by upregulating peroxisomal functions. Collectively, our findings underline the importance of metabolic plasticity in order to adapt to varying nutritional conditions to ultimately promote survival.

## 3 DISCUSSION

This study provides a detailed metabolomic and lipidomic analyses characterizing temporal effects of acute starvation on metabolite levels in both wildtype *C. elegans* and in animals lacking functional HLH-30, an ortholog of the mammalian transcription factor TFEB. As recently shown (Harvald et al., 2017), the present study reveals that starvation massively rewires metabolism and lipid regulatory networks in a manner that is strongly dependent on the transcription factor HLH-30. Here we have detected 4063 lipid species belonging to more than 10 lipid classes and show that 996 distinct lipid species in positive and 792 in negative ionization mode change upon and during starvation. Similarly, among the 2147 metabolites that we have detected, the abundance of 775 and 738 metabolites change in at least one time point across the starvation series for both wildtype and *hlh-30* animals in positive and negative ionization mode, respectively. In particular, we find that distinct cardiolipin and acyl-carnitine species change upon starvation but only in wildtype animals and not in *hlh-30* animals. Interestingly, these lipids are almost exclusively found or synthesized in mitochondria. Consistently, our findings show that both lipolysis of stored lipids and mitochondrial fatty acid oxidation are required for survival during starvation. Interestingly, supplementation of fatty acids rescues compromised survival of *hlh-30* animals during starvation. Since fatty acid supplementation does not rescue longevity of *hlh-30* animals after knockdown of the gene encoding the peroxisomal assemble factor PRX-5, we speculate that HLH-30/TFEB is required to generate and maintain functional mitochondria, and thus suggest that *hlh-30* animals rewire metabolic activities towards peroxisomal activities to sustain survival during starvation. Interestingly, HLH-30/TFEB recognizes and binds to so called CLEAR motifs (Grove et al., 2009; Lapierre et al., 2013; Settembre et al., 2013), which are found in numerous genes encoding mitochondrial proteins (Table S1). Accordingly, revisiting our previous expression analyses (Harvald et al., 2017) we find that genes encoding for mitochondrial proteins e.g. *cpt-1, acs-2* and *hacd-1* are downregulated in *hlh-30* animals compared to wildtype, while genes encoding for proteins involved in peroxisomal fatty acid metabolism e.g. acyl-CoA oxidases, are upregulated in *hlh-30* animals compared to wildtype animals (Figure S5). Our observations are perfectly aligned with recent observations by Weir et al. showing that remodeling of mitochondrial networks is required for AMPK- and dietary restriction-mediated longevity, and that lifespan depends on both fatty acid oxidation and peroxisomal functions (Weir et al., 2017). Furthermore, although HLH-30/TFEB primarily has been linked to lysosomal biogenesis, autophagy, and immune functions (Lapierre et al., 2013; Settembre et al., 2013; Visvikis et al., 2014), TFEB also exerts a global transcriptional control on glucose uptake and metabolism, fatty acid oxidation, oxidative phosphorylation, and mitochondrial biogenesis at the transcriptional level in rodents, among others via *Ppargc1α* and *Ppar1α* (Mansueto et al., 2017; Settembre et al., 2013).

The observation that fatty acid supplementation rescues lifespan of *hlh-30* animals and extends longevity of wildtype animals, suggests that mobilization of fatty acids from stored lipids is required for maintaining lifespan during starvation. Indeed, lipases like ATGL-1, LIPL-2, LIPL-3 and LIPL-5 are upregulated by dietary restriction and required for dietary restriction-induced extension of lifespan (Buis et al., 2019; Lee et al., 2014; Murphy et al., 2019; Zaarur et al., 2019). Markedly, we recently found that loss of HLH-30 diminishes expression of *atgl-1, lipl-2, lipl-3 and lipl-4* (Harvald et al., 2017). Here we found that RNAi against *atgl-1* shortens survival of wildtype animals during starvation but not of *hlh-30* animals. However, fatty acid supplementation extends lifespan of both wildtype and *hlh-30* animals, further supporting that mobilization of fatty acids from lipid stores provide metabolic energy to support organismal lifespan. Consistent with our findings, Macedo et al. recently reported that the abundance of certain cardiolipin species increases in response to dietary restriction (Macedo et al., 2019), which increased further upon loss of LIPL-5.

Collectively, this study provides a comprehensive analyses of temporal starvation responses that combines both metabolomic and lipidomic analyses. Our data highlight the relevance of combining global profiling analyses to further understand how metabolism is regulated and how an organism adapts to nutritional changes. As exemplified by our comparison of the effect of starvation on metabolites and lipids in wildtype and *hlh-30* animals, the genetic tractability of *C. elegans* and combined with RNA interference, shows that the nematode system serves as an excellent framework to delineate conserved mechanisms whereby specific metabolic pathways regulate starvation responses.

## Acknowledgements

This work was supported by The Danish Council for Independent Research, Natural Sciences (6108-00268A). We gratefully acknowledge scientific discussions with Marta Moreno-Torres.

## Author contributions

K.B.D., M.W. and N.J.F. designed experiments. K.B.D., M.W. and N.J.F. wrote the manuscript. K.B.D., E.B.H., J.F.H. and M.W. performed experiments and analyzed data.

## Data availability Statement

Data and R scripts have been deposited and are available at www.mendeley.com http://dx.doi.org/10.17632/2r27v9yjp5.1

## 4 EXPERIMENTAL PROCEDURES

### 4.1 *C. elegans* strains and maintenance

The wildtype N2 Bristol and the *hlh-30* mutant (tm1978, a kind gift from Dr. Marlene Hansen, Sandford-Burnham Medical Research Institute) were used and cultivated under standard conditions and handled as described (Harvald et al., 2017).

### 4.2 Survival-span assay

Worms were synchronized and grown until L4 stage on NGM seeded plates. For RNAi treatment worms were transferred to plates containing IPTG seeded with the respective HT115 RNAi bacteria clone. At L4, worms were transferred to empty plates to induce starvation conditions. For supplementation with fatty acids, worms were transferred to plates containing either 40 µM lauric acid (C_12:0_) or palmitic acid (C_16:0_). 12 worms were placed on each plate and scored every day. A worm was scored dead when it was unresponsive to gentle prodding.

### 4.3 Sample harvest for mass spectrometry

Worms were synchronized as described above and grown on NGM seeded plates until L4 stage and then transferred to empty NGM plates to induce starvation condition. Worms were harvested at specified time points by washing off plates with MS-grade H_2_O and washed thrice. After the final wash, worms were incubated under rotation at RT for 20 min in H_2_O. Samples were spun down for 1 min at 1,100 rpm / RT and supernatant aspirated. Samples were flash frozen in liquid nitrogen and stored at -80°C.

### 4.4 BUME extraction

Samples were extracted by using the BUME method as described (Lofgren et al., 2012) with minor modifications. Briefly, 50 µL ice-cold methanol was added to each sample and transferred to beat-beating tubes (NucleoSpin Bead Tubes Type A, Macherey Nagel, Düren, Germany). The samples were beat-beaten for three times 10 seconds with 20 seconds pause in a Precellys Beat Beating system (Bertin Technologies, Montigny-le-Bretonneux, France). The additional Cryolys module was used with liquid nitrogen to prevent excessive heating of samples during disruption. 150 µL butanol and 200 µL heptane-ethyl acetate (3:1) was added to each sample sequentially which were then incubated for 1 h at 500 rpm / RT. 200 µL 1% acetic acid was added to each sample followed by centrifugation for 15 min at 13000 rpm / 4°C. The upper organic phase was transferred to a fresh Eppendorf tube and the lower aqueous phase was re-extracted by the addition of 200 µL heptane-ethyl acetate followed by incubation and centrifugation as described above. The upper organic phase was transferred to the already obtained organic phase. The lower phase was transferred to a new Eppendorf tube and used for metabolomic analyses. Samples were evaporated to dryness and stored at -20°C. For lipidomics, samples were re-dissolved in 50 µL 65% isopropanol/ 35% acetonitrile/ 5 % H_2_O, vortexed and 40 µL were transferred to an autosampler vial. The remaining 10 µL were pooled to form a QC sample for the entire study. The precipitated proteins were used for determination of protein content using a Bicinchoninic Acid Protein Assay Kit (Sigma-Aldrich, Taufkirchen, Germany).

For metabolomics, samples were re-dissolved in 30 µL 1% formic acid, spun down for 5 min at 16000g / RT and 25 µL were transferred to HPLC vials. The remaining 5 µL were pooled to form a QC sample for the entire study.

### 4.5 Lipid analysis, data processing, and statistical analysis

Lipids were analyzed as previously described (Witting et al., 2014). Briefly, lipids were separated on a Waters Acquity UPLC (Waters, Eschborn, Germany) using a Waters Cortecs C18 column (150 mm x 2.1 mm ID, 1.6 µm particle size, Waters, Eschborn Germany) and a linear gradient from 68% eluent A (40% H_2_O / 60% acetonitrile, 10 mM ammonium formate and 0.1% formic acid) to 97% eluent B (10% acetonitrile / 90% isopropanol, 10 mM ammonium formate and 0.1% formic acid). Mass spectrometric detection was performed using a Bruker maXis UHR-ToF-MS (Bruker Daltonic, Bermen, Germany) in positive and negative ionization mode using data dependent acquisition to obtain MS^1^ and MS^2^ information. For every ten samples a pooled QC was injected to check performance of the UPLC-UHR-ToF-MS system and were used for normalization.

Raw data was processed with Genedata Expressionist for MS 12.0 (Genedata AG, Basel, Switzerland). Preprocessing steps included noise subtraction, m/z recalibration, chromatographic alignment and peak detection and grouping. Data was exported for Genedata Expressionist for MS 12.0 Analyst statistical analysis software and as .xlxs for further investigation. Maximum peak intensities were used for statistical analysis and data was normalized on the protein content of the sample and an intensity drift normalization based on QC samples was used to normalize for the acquisition sequence.

Lipids were putatively annotated on the MS^1^ level using an in-house developed database for *C. elegans* lipids and bulk composition from LipidMaps (O’Donnell, Dennis, Wakelam, & Subramaniam, 2019), when available. MS2 data was extracted using Bruker Data Analysis 5.0 (Bruker Daltonics) and read into R using the MSnbase package (Gatto & Lilley, 2012). Matching against an *in-silico* database of *C. elegans* lipids and LipidBlast was performed using the masstrixR package (Kind et al., 2013) (Witting unpublished, https://github.com/michaelwitting/masstrixR) and only hits with a forward and reverse matching score > 0.75 were considered. Lipids with a *p*-value < 0.05 and a foldchange of > 2 or < 0.5 were considered to be significantly changed. Annotations of interesting biological peaks were manually verified and corrected if necessary.

### 4.6 Metabolomics analysis and data processing of BUME extraction

5 µL were injected using a Vanquish Horizon UPLC (Thermo Fisher Scientific, Germering, Germany) and compounds separated on a Zorbax Eclipse Plus C18 guard (2.1 × 50 mm and 1.8 μm particle size, Agilent Technologies, Santa Clara, CA, USA) and an analytical column (2.1 × 150 mm and 1.8 μm particle size, Agilent Technologies, Santa Clara, CA, USA) kept at 40°C. The analytes were eluted using a flow rate of 400 μL/min and the following composition of eluent A (0.1% formic acid) and eluent B (0.1% formic acid, acetonitrile) solvents: 3% B from 0 to 1.5 min, 3–40% B from 1.5 to 4.5 min, 40-95% B from 4.5 to 7.5 min, 95 % B from 7.5 to 10.1 min and 95 to 3% B from 10.1 to 10.5 min before equilibration for 3.5 min with the initial conditions. Using a 6-port valve and a secondary pump, the guard wash backflushed from 9-10 min with a flow of 1 mL/min with 95% B. The flow from the UPLC was coupled to a Q Exactive HF mass spectrometer (Thermo Fisher Scientific, Bremen, Germany) for mass spectrometric analysis in both positive and negative ion mode using the following general settings for MS1 mode: resolution: 120,000, AGC target: 3e6, maximum injection time: 200 ms, scan range 65-975 m/z and lock mass: 391.28429/112.98563 (pos/neg mode). For compound fragmentation MS1/ddMS2 mode were used with the following general settings: resolution: 60,000/15,000, AGC target: 1e6/1e5, maximum injection time: 50/100 ms, scan range 65-975 m/z, loop count: 5, isolation with: 2 m/z and normalized collision energy: 35/38 (pos/neg). The generated pooled sample was used for quality control (QC) and compound fragmentation. Samples were analyzed in randomized order with a QC sample injected every 9^th^ run. A MS2 inclusion list was generated from the blanks and the last equilibration runs using Compound Discover v. 3.0 (Thermo Fisher Scientific) to ensure fragmentation of the later extracted features. The features found in the blank were removed from the inclusion list if they were not > 5 x more abundant in the QC samples.

Raw data was processed with MzMine (v 2.42) (Pluskal, Castillo, Villar-Briones, & Oresic, 2010). In brief, the following modules were used: Mass detection, ADAP chromatogram builder, ADAP deconvolution, Join aligner, Isotopic peak grouper, Gap filling (same RT and m/z range) and Identification in local spectra database search; all with 5 ppm mass tolerance and 0.25 RT tolerance when possible. Final peaklist included features found in at least 60% of the samples, which had at least 2 peaks in an isotope pattern. Compounds were annotated at Metabolomics Standards Initiative (MSI) (Salek, Haug, & Steinbeck, 2013) level 2 using local MS/MS spectra databases of National Institute of Standards and Technology 17 (NIST17) and MassBank of North America (MoNA). Moreover, the MS/MS data were also searched in MzCloud using Compound Discoverer for additional annotations at MSI level 2. MSI level 3 annotation were achieved using SIRIUS (Duhrkop et al., 2019) before lastly MSI level 4 annotation by searching in Human Metabolite Database (Wishart et al., 2018). After compound annotation, the datasets were corrected for signal drift using the R package statTarget (Luan et al., 2018). Metabolites with a *p*-value < 0.05 and a foldchange of > 2 or < 0.5 were considered to be significantly changed.

### 4.7 MeOH/ACN/H_2_O extraction of metabolites

Samples were harvested at described above (2000 worms pr. sample in biological triplicates) and flash frozen to be stored at -80 °C. Samples were thawed on ice and 200 µL ice-cold extraction solvent (50% methanol/ 30% acetonitrile / 20% H_2_O, MS-grade) was added to each sample followed by vigorous vortexing. Samples were sonicated in an ice bath for 10 × 30 sec at high speed. Samples were incubated in a Thermoshaker for 30 min at 1.200 rpm / 4°C, vortexed vigorously and spun down for 10 min at 12,000g / 4°C. The supernatant was transferred to fresh Eppendorf tubes, lyophilized and stored at -20°C. Before injection samples were re-dissolved in 30 µL 0.1 % formic acid, spun down for 5 min at 16,000g / RT and supernatant transferred to HPLC vials.

### 4.8 Metabolomics analysis and data processing of MeOH/ACN/H_2_O extracted metabolites

Metabolites were analyzed according to (Sustarsic et al., 2018) with minor alterations to the gradient. In brief, metabolites were analyzed with LC-MS using reverse phase (RP) separation. 5 μL were injected using an Agilent 1290 Infinity HPLC system (Agilent Technologies, Santa Clara, CA, USA) equipped with an Agilent Zorbax Eclipse Plus C18 column (2.1 × 150 mm, 1.8 μm, Agilent Technologies, Santa Clara, CA, USA) with a 50 mm guard-column, both kept at 40°C. The analytes were eluted using a flow rate of 300 µL/min and solvent with the following eluent composition: eluent A (0.1% formic acid, H_2_O) and eluent B (0.1% formic acid, acetonitrile): 97% A from 0-8 min, 60% A from 8-12 min, 10% A from 12-15 min before equilibration with initial conditions for 3 min. The HPLC flow was coupled to an Agilent 6530 quadrupole time of flight (Q-TOF) mass spectrometer scanning from 70-1000 m/z. A pooled sample was generated for compound fragmentation and all samples were analyzed in both positive and negative ion mode. Samples were run in all-ion fragmentation mode with collision energy of 20 V, in order to produce fragments for identification of metabolites. Libraries of metabolites with retention time (RT) were constructed using Agilent MassHunter PCDL Manager. The identification of each compound was based on exact mass, RT and/or comparison of fragments with the Metlin MS/MS database (https://metlin.scripps.edu). Chromatograms for all compounds were extracted and quantified using Agilent Profinder using a mass tolerance of 20 ppm and a retention time tolerance of 0.1 min.

### 4.9 β-oxidation assay

The method for measuring fatty acid oxidation was applied as described previously (Elle, Rodkaer, Fredens, & Faergeman, 2012).

