## Supplementary material for "HLH-30 dependent rewiring of metabolism during starvation in *C. elegans*": Suppl. info

**Supplemental material**
**Dall et al.**

Figure S1

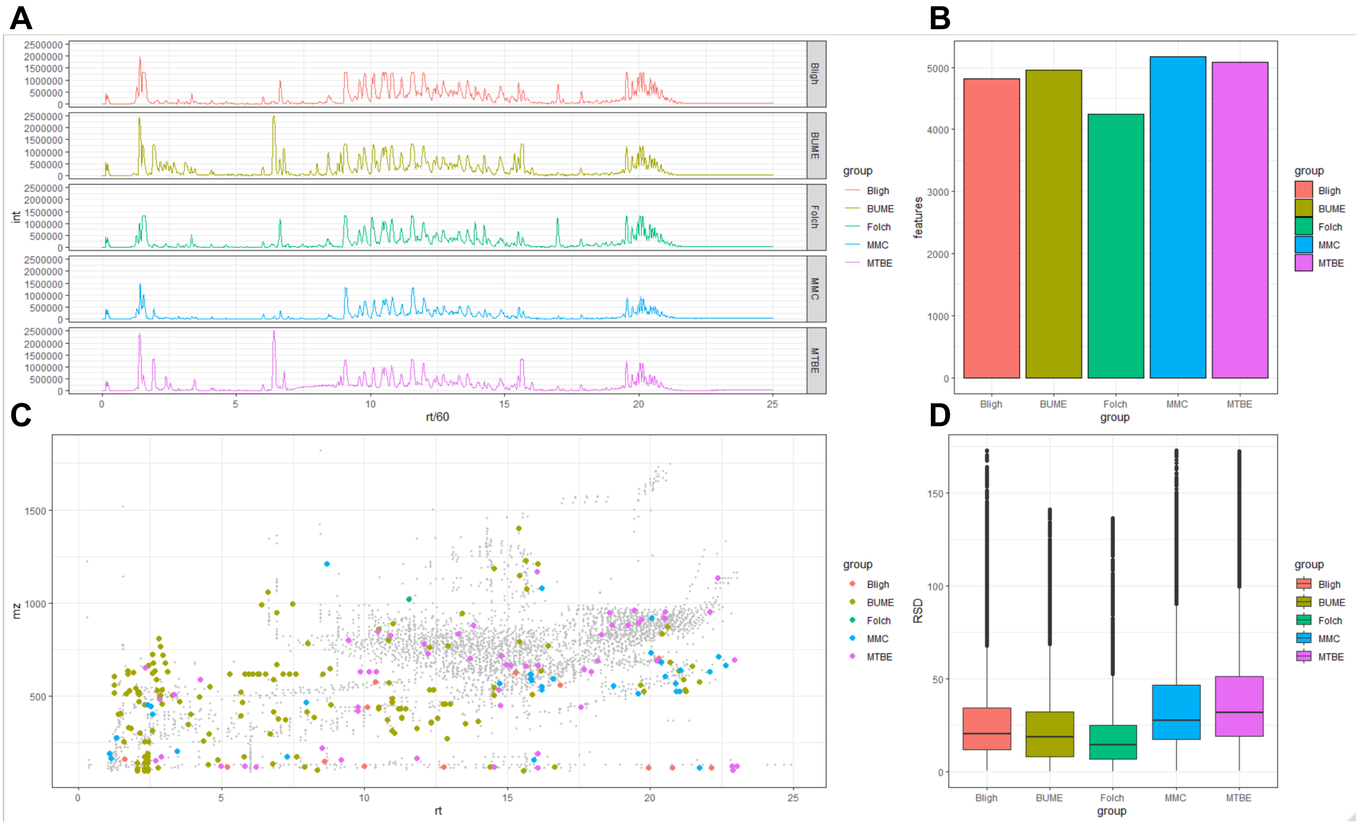


**Figure S1. Comparison of lipid extraction methods.**
(A) Base peak chromatogram for each of the extraction methods: Bligh and Dyer (red), BUME (yellow), Folch (green), MMC (blue) and MTBE (purple).
(B) Bar plot showing the number of features detected for each extraction method.
(C) Scatter plot showing m/z values and retention time (RT) for the features of each extraction method. Shown in grey are the common features for all the extraction methods.
(D) Boxplot showing the relative standard deviation (RSD) between the replicates of each extraction method.

Figure S2

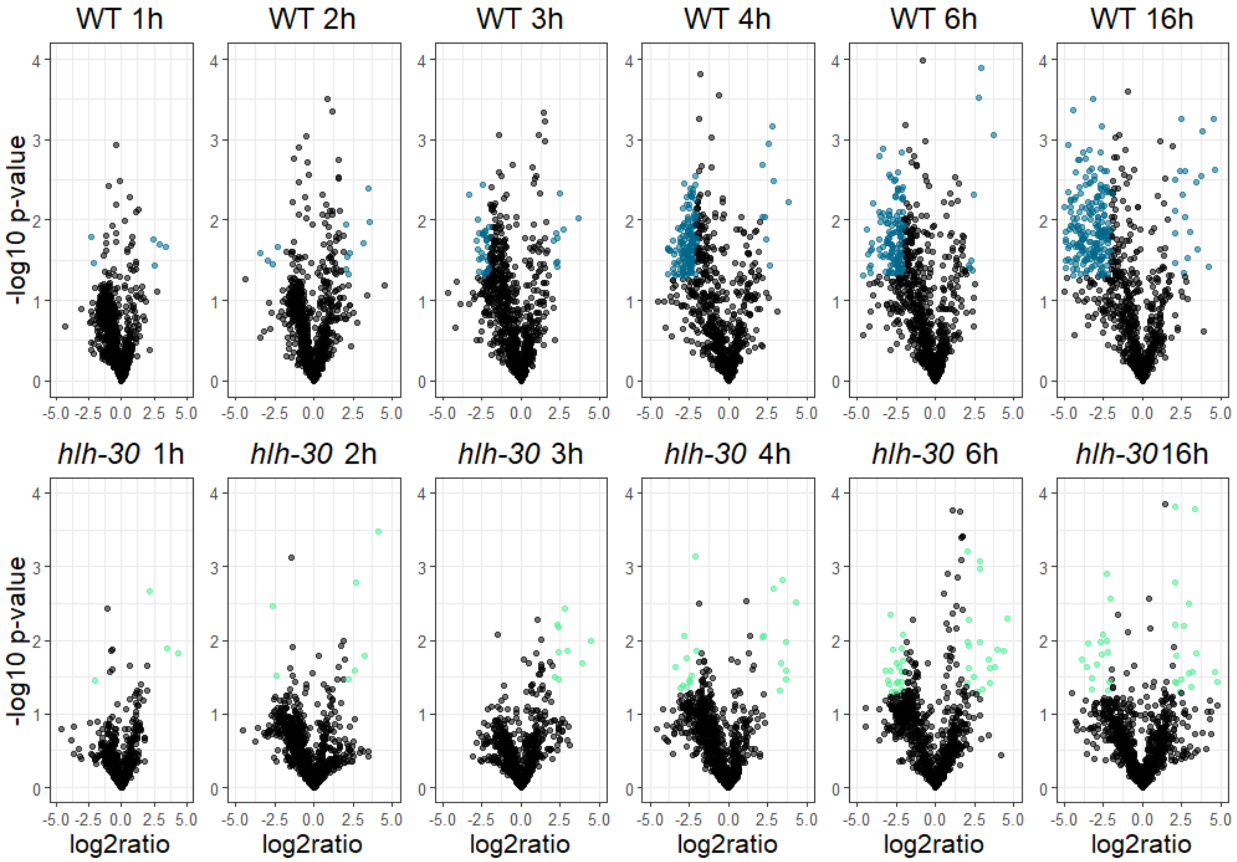


**Figure S2 related to Figure 2A. Metabolic changes induced by temporal starvation.**

Volcano plots displaying changes in the metabolome in response to starvation identified in negative ionization mode. Significant up- or down-regulated metabolites in the wildtype are shown in blue, and in light green for the *hlh-30* mutant.

Figure S3

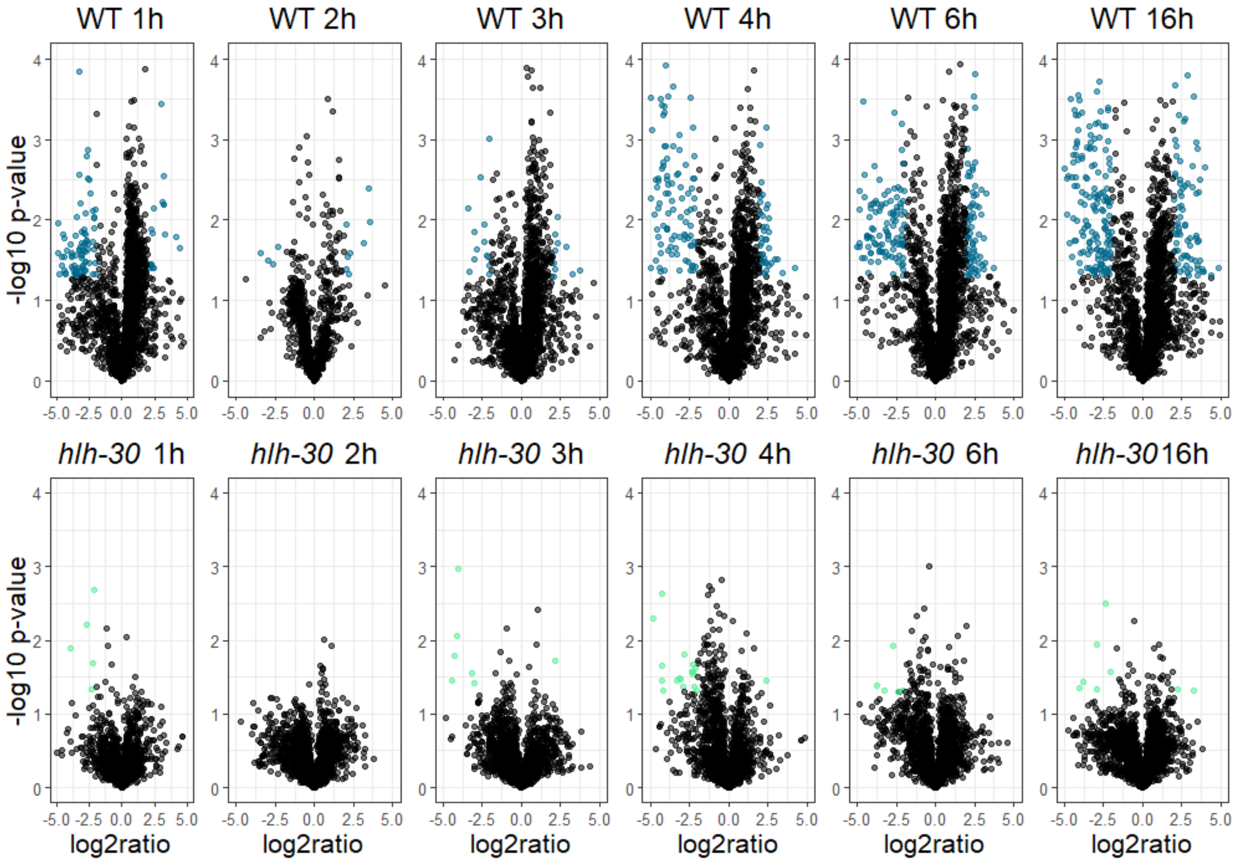


**Figure S3 related to Figure 2B. Lipidomic changes induced by temporal starvation.**

Volcano plot displaying changes in the lipidome in response to starvation identified in negative ionization mode. Significant up- or down-regulated lipids in the wildtype are shown in blue, and in light green for the *hlh-30* mutant.

Figure S4

**A**

| mz | RT | anno | mz (theo.) | error (abs) | error (ppm) |
|---|---|---|---|---|---|
| 1539.0041 | 17.9 | CL(78:14) | 1539.0060 | 0.0019 | 1.2 |
| 1541.0203 | 18.2 | CL(78:13) | 1541.0217 | 0.0014 | 0.9 |
| 1543.0341 | 18.6 | CL(78:12) | 1543.0373 | 0.0032 | 2.1 |
| 1563.0049 | 17.5 | CL(80:16) | 1563.0060 | 0.0011 | 0.7 |
| 1565.0206 | 17.9 | CL(80:15) | 1565.0217 | 0.0011 | 0.7 |
| 1567.0382 | 18.3 | CL(80:14) | 1567.0374 | -0.0009 | -0.6 |
| 1569.0508 | 18.7 | CL(80:13) | 1569.0530 | 0.0022 | 1.4 |

**B**

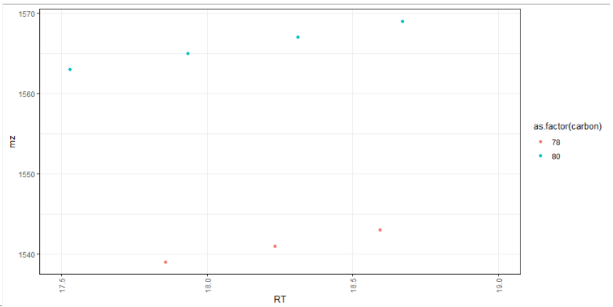


**C**

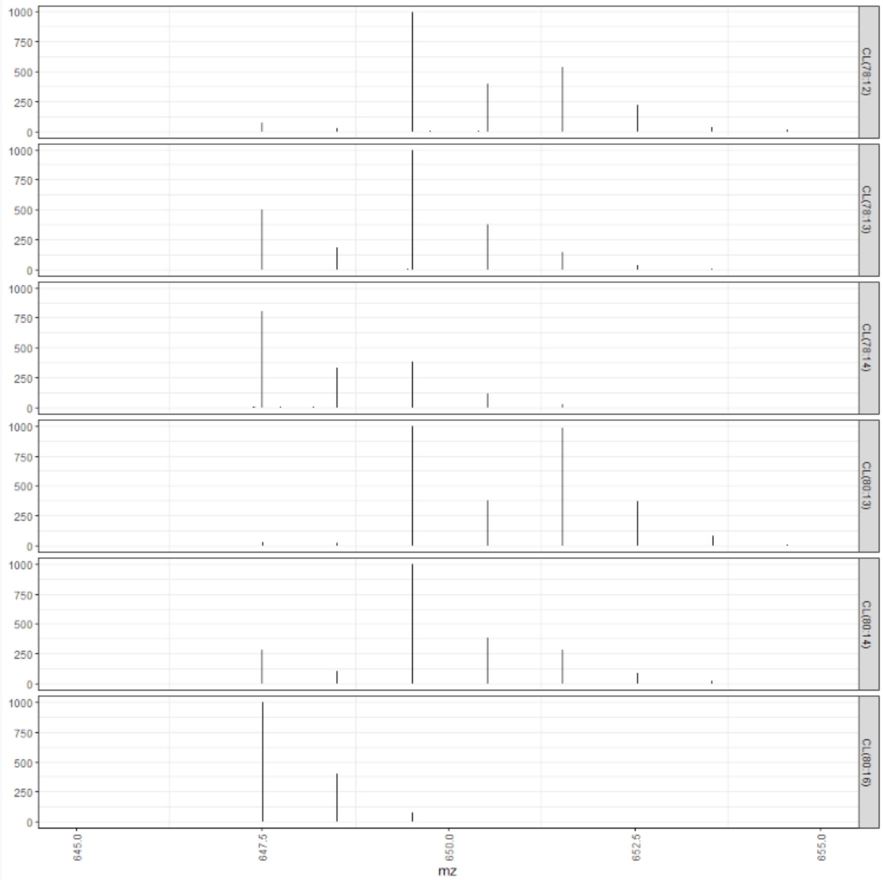


**Figure S4. Details on cardiolipin annotation.**

(A) Annotation of MS1 data with different cardiolipins (CL) detected as $[M+NH_4]^+$ adduct ion.

(B) RT trend plots for the detected CLs. Trends along the retention time dimension indicate a homologous series of lipids belonging to the same lipid class. The groups are colored according to the number of carbons in the acyls.

(C)For all six of the seven putative cardiolipins MS/MS spectra were available. CLs were detected in positive ion mode. No $[M-H]^-$ or $[M-2H]2^-$ peaks were detected in negative ion mode due to generally lower sensitivity in negative ion mode. The fragmentation pattern showed prominent fragments, which were equal to a $[M+H-H_2O]^+$ of DGs. Therefore, a neutral loss of the phosphorylated glycerol backbone with two attached fatty acids in combination with the $NH_4$ occurs. Therefore, “symmetric” cardiolipins would show only one specific fragment, while asymmetric ones have different fragments. The fragmentation pattern of the lipid feature annotated as CL(80:16) showed only one fragment corresponding to DG(40:8) $[M+H-H_2O]^+$. In contrast the cluster annotated as CL(80:14) showed three different peaks representing DG(40:8), DG(40:7), DG(40:6). Using this information two possibilities to form CL(80:14) exist and peaks are co-eluting. Based on the fragment intensities the symmetric CL is the major species. Fragments of all other lipid clusters were in agreement with the described fragmentation.

Figure S5

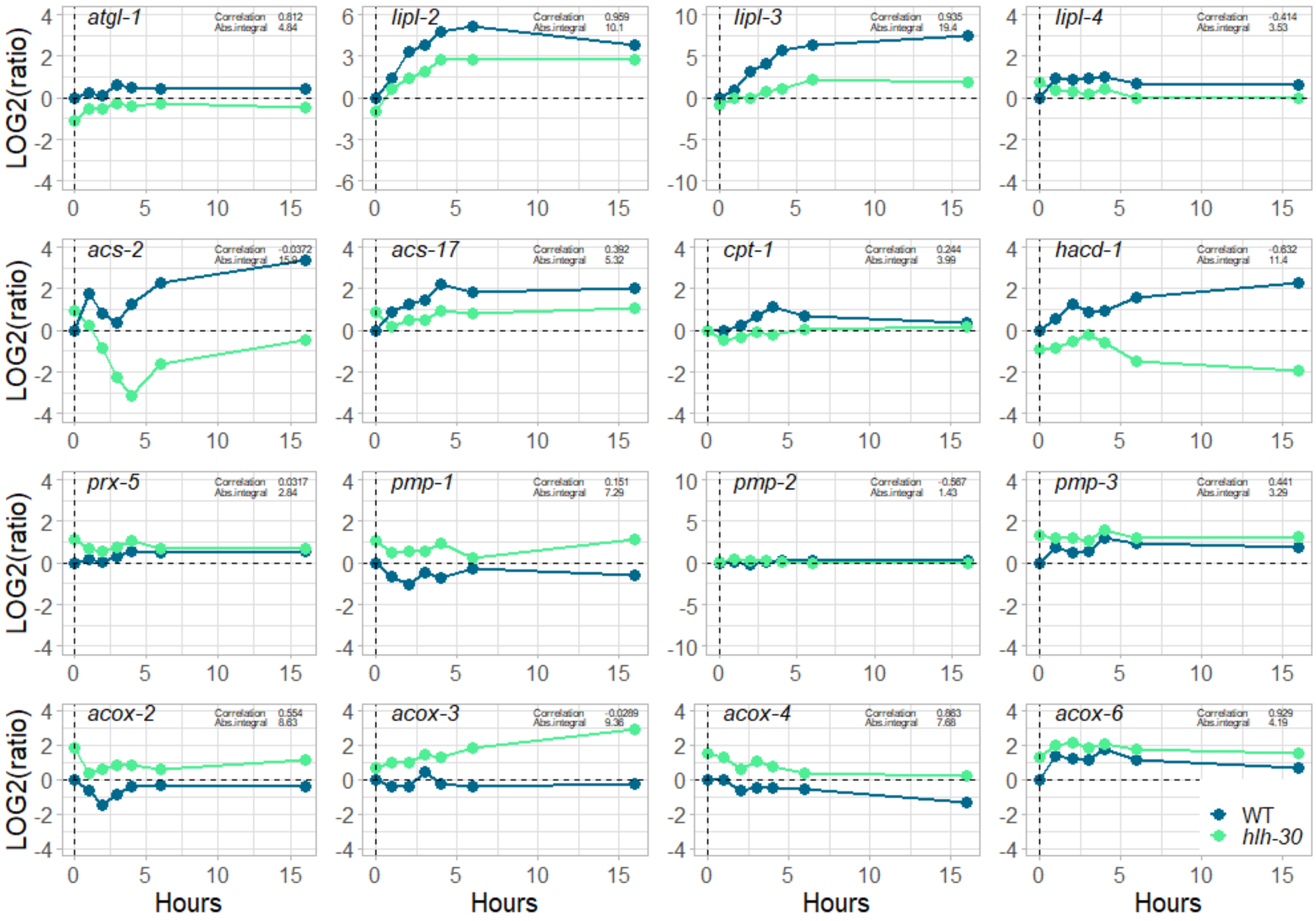


**Figure S5. Expression levels of selected genes determined with RNA-seq in *C. elegans*.** Line plots displaying changes in gene expression of lipid catabolism genes during the starvation timeseries in wildtype (blue) and the *hlh-30* mutant (light green). Lipolytic lipases *atgl-1*, *lipl-2*, *lipl-3* and *lipl-4* show decreased expression in *hlh-30* mutant when compared to wildtype animals. Fatty acyl activating genes *acs-2* and *acs-17* are likewise decreased in the *hlh-30* mutant in addition to mitochondrial β-oxidation genes *cpt-1* and *hacd-1*. Peroxisomal assembly factor and fatty acyl transporter, *prx-5* and *pmp-2* respectively show little change in expression in response to starvation while two other fatty acyl transporter genes *pmp-1* and *pmp-3* show increase in expression for the *hlh-30* mutant when compared to wildtype animals. Peroxisomal β-oxidation genes, *acox-2*, *acox-3*, *acox-4* and *acox-6* all show increased expression during starvation in the *hlh-30* mutant when compared to wildtype levels. See also Harvald et al. (2017) for details.

### Figures and Tables

| Strain | FA treat-ment | FA treated survival -span (d) | FA treated survival-span (d)* | No. of treated animals | Control survival-span (d) | Control survival-span (d)* | No. of Control animals | Max survival-span (d) | *p*-value versus Control |
|---|---|---|---|---|---|---|---|---|---|
| **N2 Bristol** (WT) | $C_{12:0}$ | 11 (45.4%) | 10.75 (36.5%) | 93/120 | 6 | 6.82 | 91/120 | 16/10 | <0.0001 |
| | $C_{12:0}$ | 10 (20%) | 9.96 (26.9%) | 85/120 | 8 | 7.28 | 93/120 | 17/10 | <0.0001 |
| | $C_{12:0}$ | 13 (53.8%) | 12.03 (49.6%) | 95/120 | 6 | 6.06 | 93/120 | 16/10 | <0.0001 |
| | $C_{16:0}$ | 8 (25%) | 7.74 (21.7%) | 109/120 | 6 | 6.06 | 93/120 | 12/10 | <0.0001 |
| | $C_{16:0}$ | 9 (11.1%) | 8.53 (13.0%) | 94/120 | 8 | 7.42 | 97/120 | 12/10 | <0.0001 |
| | $C_{16:0}$ | 9 (11.1%) | 7.89 (5.70%) | 106/120 | 8 | 7.44 | 87/120 | 12/10 | 0.0045 |
| **JIN1375** (*hlh-30*) | $C_{12:0}$ | 10 (40%) | 8.79 (37.8%) | 103/120 | 6 | 5.46 | 81/120 | 15/8 | <0.0001 |
| | $C_{12:0}$ | 9 (44.4%) | 9.43 (46.0%) | 103/120 | 5 | 5.09 | 89/120 | 16/7 | <0.0001 |
| | $C_{12:0}$ | 13 (61.5%) | 10.78 (62.2%) | 104/120 | 5 | 4.07 | 88/120 | 15/7 | <0.0001 |
| | $C_{16:0}$ | 8 (37.5%) | 7.43 (45.2%) | 98/120 | 5 | 4.07 | 88/120 | 9/7 | <0.0001 |
| | $C_{16:0}$ | 8 (50%) | 7.51 (43.6%) | 98/120 | 4 | 4.23 | 90/120 | 10/6 | <0.0001 |
| | $C_{16:0}$ | 9 (44.4%) | 7.29 (37.4%) | 102/120 | 5 | 4.56 | 89/120 | 10/7 | <0.0001 |

**Supplementary Table S2. Summary of survival-span statistics.** Median, mean and maximum survival and statistical significance were calculated for each experiment. *Mean survival lifespan. *p*-values were based on median survival lifespan.

| Strain | RNAi treat-ment | FA treat-ment | RNAi / FA treated survival-span (d) | RNAi / FA treated survival-span (d)* | No. of treated animals | Control survival-span (d) | Control survival-span (d)* | No. of Control animals | Max survival -span (d) | *p*-value versus Control |
|---|---|---|---|---|---|---|---|---|---|---|
| **N2 Bristol** (WT) | *prx-5* (JA) | none | 8 (0%) | 7 (-3.28%) | 89/120 | 8 | 7.23 | 89/120 | 11/10 | 0.4624 |
| | *prx-5* (JA) | none | 8 (-12.5%) | 7.36 (-2.58%) | 88/120 | 9 | 7.55 | 83/120 | 10/10 | 0.2838 |
| | *prx-5* (JA) | none | 9 (11.11%) | 7 (-1%) | 85/120 | 8 | 7.07 | 84/120 | 10/10 | 0.7177 |
| | *atgl-1* (JA) | none | 6 (-33.32%) | 5.21 (-38.77%) | 88/120 | 8 | 7.23 | 89/120 | 8/10 | <0.0001 |
| | *atgl-1* (JA) | none | 7 (-28.57%) | 6.47 (-16.69%) | 88/120 | 9 | 7.55 | 83/120 | 8/10 | < 0.0001 |
| | *atgl-1* (JA) | none | 5 (-60%) | 4.75 (-48.84%) | 84/120 | 8 | 7.07 | 84/120 | 8/10 | < 0.0001 |
| | *cpt-1* (JA) | none | 3 (-166.66%) | 3.38 (-106.21%) | 83/120 | 8 | 6.97 | 86/120 | 5/10 | < 0.0001 |
| | *prx-5* (JA) | $C_{12:0}$ | 12 (0%) | 9.92 (-6.04%) | 88/120 | 12 | 10.52 | 87/120 | 15/15 | 0.3020 |
| | *prx-5* (JA) | $C_{12:0}$ | 12 (8.33%) | 10.17 (-3.34%) | 84/120 | 11 | 10.51 | 86/120 | 15/15 | 0.8463 |
| | *prx-5* (JA) | $C_{12:0}$ | 10 (-10%) | 9.46 (-2.21%) | 105/120 | 11 | 9.67 | 107/120 | 15/15 | 0.2236 |
| | *atgl-1* (JA) | $C_{12:0}$ | 11 (-9.09%) | 9.57 (-9.92%) | 89/120 | 12 | 10.52 | 87/120 | 15/15 | 0.1780 |
| | *atgl-1* (JA) | $C_{12:0}$ | 12 (8.33%) | 10.59 (0.75%) | 88/120 | 11 | 10.51 | 86/120 | 15/15 | 0.8616 |
| | *atgl-1* (JA) | $C_{12:0}$ | 12 (8.33%) | 9.76 (0.92%) | 95/120 | 11 | 9.67 | 107/120 | 15/15 | 0.9674 |
| | *cpt-1* (JA) | $C_{12:0}$ | 9 (-11.11%) | 8.11 (-14.05%) | 102/120 | 10 | 9.25 | 98/120 | 15/15 | 0.2773 |
| | *prx-5* (JA) | $C_{16:0}$ | 8.5 (-41.17%) | 8.61 (-12.07%) | 104/120 | 12 | 9.65 | 89/120 | 14/14 | 0.0228 |
| | *prx-5* (JA) | $C_{16:0}$ | 9 (-22.22%) | 9.17 (-4.14%) | 91/120 | 11 | 9.55 | 85/120 | 14/14 | 0.462 |
| | *prx-5* (JA) | $C_{16:0}$ | 10 (-5%) | 9 (-2.77%) | 101/120 | 10.5 | 9.25 | 110/120 | 14/14 | 0.3309 |
| | *atgl-1* (JA) | $C_{16:0}$ | 11 (-9.09%) | 9.78 (1.32%) | 87/120 | 12 | 9.65 | 89/120 | 14/14 | 0.5313 |
| | *atgl-1* (JA) | $C_{16:0}$ | 12 (8.33%) | 10.18 (6.18%) | 89/120 | 11 | 9.55 | 85/120 | 14/14 | 0.1805 |
| | *atgl-1* (JA) | $C_{16:0}$ | 11.5 (8.33%) | 9.38 (1.38%) | 102/120 | 10.5 | 9.25 | 110/120 | 14/14 | 0.6918 |
| | *cpt-1* (JA) | $C_{16:0}$ | 3 (-200%) | 5.48 (-53.28%) | 104/120 | 9 | 8.4 | 97/120 | 12/15 | < 0.0001 |

**Supplementary Table S3A. Summary of survival-span statistics for wildtype animals.** Median, mean and maximum survival and statistical significance were calculated for each experiment. *Mean survival lifespan. *p*-values were based on median survival lifespan.

| Strain | RNAi treat ment | FA treat-ment | RNAi / FA treated survival-span (d) | RNAi / FA treated survival-span (d)* | No. of treated animals | Control survival-span (d) | Control survival-span (d)* | No. of Control animals | Max survival -span (d) | *p*-value versus Control |
|---|---|---|---|---|---|---|---|---|---|---|
| **JIN1375** (*hlh-30*) | *prx-5* (JA) | none | 3 (-33.33%) | 3.16 (-25.63%) | 86/120 | 4 | 3.97 | 85/120 | 6/6 | < 0.0001 |
| | *prx-5* (JA) | none | 3 (-33.33%) | 3.09 (-22.33%) | 93/120 | 4 | 3.78 | 88/120 | 5/6 | < 0.0001 |
| | *prx-5* (JA) | none | 3 (-33.33%) | 2.98 (-25.50%) | 84/120 | 4 | 3.74 | 85/120 | 5/6 | < 0.0001 |
| | *atgl-1* (JA) | none | 4 (0%) | 3.56 (-11.51%) | 90/120 | 4 | 3.97 | 85/120 | 6/6 | 0.0032 |
| | *atgl-1* (JA) | none | 4 (0%) | 3.5 (-8%) | 94/120 | 4 | 3.78 | 88/120 | 5/6 | 0.0018 |
| | *atgl-1* (JA) | none | 3 (-33.33%) | 3.15 (-18.73%) | 83/120 | 4 | 3.74 | 85/120 | 5/6 | 0.0004 |
| | *cpt-1* (JA) | none | 4 (0%) | 3.74 (3.20%) | 87/120 | 4 | 3.62 | 85/120 | 6/6 | 0.6441 |
| | *prx-5* (JA) | $C_{12:0}$ | 4 (75%) | 4.06 (-88.91%) | 87/120 | 7 | 7.67 | 84/120 | 6/14 | < 0.0001 |
| | *prx-5* (JA) | $C_{12:0}$ | 3 (-233.33%) | 3.3 (-173.63%) | 86/120 | 10 | 9.03 | 91/120 | 5/14 | < 0.0001 |
| | *prx-5* (JA) | $C_{12:0}$ | 3 (-233.33%) | 3.1 (184.83%) | 94/120 | 10 | 8.83 | 112/120 | 5/15 | < 0.0001 |
| | *atgl-1* (JA) | $C_{12:0}$ | 6 (-16.66%) | 5.5 (-39.45%) | 88/120 | 7 | 7.67 | 84/120 | 10/14 | < 0.0001 |
| | *atgl-1* (JA) | $C_{12:0}$ | 9 (-11.11%) | 7.29 (-23.86%) | 87/120 | 10 | 9.03 | 91/120 | 10/14 | < 0.0001 |
| | *atgl-1* (JA) | $C_{12:0}$ | 11.5 (13.04%) | 9.33 (5.35%) | 98/120 | 10 | 8.83 | 112/120 | 14/15 | 0.4269 |
| | *cpt-1* (JA) | $C_{12:0}$ | 9 (0%) | 8.37 (0%) | 101/120 | 9 | 8.37 | 100/120 | 15/15 | 0.4095 |
| | *prx-5* (JA) | $C_{16:0}$ | 4 (-75%) | 4.33 (-51.27%) | 100/120 | 7 | 6.55 | 90/120 | 6/10 | < 0.0001 |
| | *prx-5* (JA) | $C_{16:0}$ | 3 (-200%) | 3.19 (-118.49%) | 84/120 | 9 | 6.97 | 86/120 | 5/10 | < 0.0001 |
| | *prx-5* (JA) | $C_{16:0}$ | 3 (-200%) | 2.9 (-189.31%) | 102/120 | 9 | 8.39 | 103/120 | 5/12 | < 0.0001 |
| | *atgl-1* (JA) | $C_{16:0}$ | 6 (-16.6%) | 6.03 (-8.62%) | 91/120 | 7 | 6.55 | 90/120 | 10/11 | 0.0068 |
| | *atgl-1* (JA) | $C_{16:0}$ | 9 (0%) | 7.34 (5.04%) | 89/120 | 9 | 6.97 | 86/120 | 10/10 | 0.4547 |
| | *atgl-1* (JA) | $C_{16:0}$ | 11 (18.18%) | 8.48 (1.06%) | 94/120 | 9 | 8.39 | 103/120 | 13/14 | 0.1734 |
| | *cpt-1* (JA) | $C_{16:0}$ | 6.5 (-7.69%) | 6.05 (-3.63%) | 110/120 | 7 | 6.27 | 105/120 | 10/10 | 0.0637 |

**Supplementary Table S3B. Summary of survival-span statistics for *hlh-30* mutant animals.** Median, mean and maximum survival and statistical significance were calculated for each experiment. *Mean survival lifespan. *p*-values were based on median survival lifespan.
